# Facultative adjustment of paternal care in the face of female infidelity in dunnocks

**DOI:** 10.1101/158816

**Authors:** Eduardo S. A. Santos, Shinichi Nakagawa

## Abstract

A much-debated issue is whether or not males should reduce parental care when they lose paternity (i.e. the certainty of paternity hypothesis). While there is general support for this relationship across species, within-population evidence is still contentious. Among the main reasons behind such problem is the confusion discerning between-from within-individual patterns. Here, we tested this hypothesis empirically by investigating the parental care of male dunnocks *(Prunella modularis)* in relation to paternity. We used a thorough dataset of observations in a wild population, genetic parentage, and a within-subject centring statistical approach to disentangle paternal care adjustment within-male and between males. We found support for the certainty of paternity hypothesis, as there was evidence for within-male adjustment in paternal care when socially monogamous males lost paternity to extra-pair sires. There was little evidence of a between-male effect overall. Our findings show that monogamous males adjust paternal care when paired to the same female partner. We also show that – in monogamous broods – the proportion of provisioning visits made by males yields fitness benefits in terms of fledging success. Our results suggest that socially monogamous females that engage in extra-pair behaviour may suffer fitness costs, as their partners’ reduction in paternal care can negatively affect fledging success.

## Introduction

Parental investment is stated as an investment by the parent in the young that increases the young’s chance of survival at a cost to the parent’s future ability to invest in other offspring (Trivers 1972; Smiseth et al. 2012). Thus, an individual should only invest into providing parental care when the fitness benefits are greater than the costs. The amount of care that a parent should provide is constrained by a series of factors, but mostly by factors that affect the reproductive value of current young, and factors that affect the residual reproductive value of parents (Klug et al. 2012). From a male parent perspective, the reproductive value of a brood should depend on the benefit that the offspring will gain from his paternal care conditional on the relatedness between the offspring and himself; a relationship known as Hamilton’s rule (Hamilton 1964a; Hamilton 1964b). However, the male’s relatedness to offspring in a brood is frequently uncertain, as females often produce extra-pair offspring. Extra-pair paternity is a common phenomenon across animals (e.g., reptiles: reviewed by Uller and Olsson 2008; fish: reviewed by Coleman and Jones 2011, and it has been extensively investigated in birds, reviewed by Griffith et al. 2002). Given the costs that parental care should exert (Royle et al. 2012), the amount of paternal care should be related to a male's certainty of paternity (Trivers 1972). This intuitive relationship has received large theoretical and empirical attention (reviewed in Sheldon 2002; Griffin et al. 2013; Forstmeier et al. 2014; Schroeder et al. 2016), but the results have been ambiguous.

Different types of theoretical models have three main predictions concerning the intra-specific question of whether males should adjust paternal care when paternity is uncertain. First, an initial set of models predicted that loss of paternity may seldom or rarely influence paternal care (Maynard Smith 1978; Grafen 1980). These early models assumed that there is no paternity evaluation, and the only cost of paternal care is time-out from the mating pool. A second group of models predicted that males should decrease their total paternal care towards a brood when certainty of paternity is low (Xia 1992; Whittingham et al. 1992; Westneat and Sherman 1993). These models assumed that males have the capacity of assessing their paternity. The third and final group of models predicted that loss of paternity may not only reduce paternal care, but it may also lead to discrimination against non-kin, which would incur in changes in allocation towards individual offspring (Westneat and Sherman 1993; Johnstone 1997).

Numerous empirical studies have tested whether a negative relationship between paternal care and loss of paternity exists (hereafter, the certainty of paternity hypothesis) (reviewed in Griffin et al. 2013). While there is general meta-analytic support for the certainty of paternity hypothesis (Griffin et al. 2013), there is also unexplained variation that may be caused by the sampling design of the study. The discussions on what approaches are most appropriate to examine the certainty of paternity hypothesis were published almost 20 years ago (Kempenaers and Sheldon 1997; Kempenaers and Sheldon 1998; Lifjeld et al. 1998; Wagner et al. 1998; Sheldon 2002), but are still pertinent given the large variation reported recently (Griffin et al. 2013). Kempenaers and Sheldon (1997), and Sheldon (2002) suggested that in order to reduce the effect of confounding factors, and to appropriately test the certainty of paternity hypothesis, two approaches should be used: 1) the analysis of sequential breeding attempts by the same male-female pair, or 2) experimental studies that remove/detain mated males or females during the fertile period of the female. Almost certainly due to logistic difficulties of the first approach, most studies to date have examined the certainty of paternity hypothesis either using simple correlational or experimental evidence across broods from different pairs (see Griffin et al. 2013 for a recent meta-analysis that includes both types of studies; and see Schroeder et al. 2016 for an example of the first approach).

The aim of this study was to test whether males reduce their paternal care (i.e. certainty of paternity hypothesis) or not when their share of paternity decreases in a brood. To achieve this aim, we conducted a field study that replicated the seminal investigation by Burke et al. (1989) of paternity and paternal care in the dunnock, *Prunella modularis,* using a population introduced into New Zealand. Dunnocks breed in ‘cooperatively’ polyandrous groups, and females copulate with several males; both their social partners and extra-group males (Burke et al. 1989; Santos et al. 2015a). It has been shown (Davies 1992) that male dunnocks provide paternal care in relation to their share of mating access with a female. More specifically, this only occurred in polyandrous and polygynandrous groups when a second male also fed the young (Davies 1992). Thus, male dunnocks are likely to have been selected to perceive their certainty of paternity. We investigated whether variation in paternal care (within-male and between-males, and within-pair and between-pairs) is associated with levels of paternity using a simple statistical approach, within-subject centring (see van de Pol and Wright 2009). We also estimated the repeatability of paternal care as a further test of whether males alter their levels of care between different breeding attempts in which paternity often differed. Moreover, we investigated whether levels of paternal care yielded benefits to current offspring or costs to the males that provided care.

## Materials and methods

### General procedures

Dunnocks have a complex and variable social mating system, with birds from the same population breeding in socially monogamous, polyandrous, polygynous or polygynandrous groups (Davies 1992; Santos and Nakagawa 2013). We studied a population of dunnocks in a 7-hectare area of the Dunedin Botanic Garden, Dunedin, New Zealand, during three breeding seasons (September to January, 2009-2012). Dunnocks were introduced into New Zealand in the 19^th^ century, and approximately 185 individuals were released around Dunedin (see Santos 2012; Moulton et al. 2014, but see Pipek et al. 2015). Effective population size at our study site has been estimated at 54 breeding individuals (Santos et al. 2013). All adults and nestlings were individually colour ringed (nestlings: day 9), and blood sampled under the Animal Ethics Committee of the University of Otago, permit no. 08/09 and the New Zealand National Bird Banding Scheme Institutional Permit to Band Birds no. 2008/075.

### Genotyping and assignment of paternity

To determine paternity, we screened the genotypes of individuals using a set of 16 polymorphic microsatellite loci (for further details of molecular methods see Santos et al. 2015a, and Santos et al. 2015b). We assigned paternity using *MasterBayes* version 2.47 (Hadfield et al. 2006) (see Santos et al. 2015a for details). Overall, irrespective of social group composition*,MasterBayes* assigned 17.0% (49/288) of chicks as extra-group young (EGY), and 26.5% of broods (26/98) contained ≥ 1 EGY. Of the chicks of socially monogamous pairs, 23.3% (32/137) were assigned as EGY, whereas only 9.9% (14/143) of the chicks of socially polyandrous groups were assigned as EGY. Within polyandrous groups, subordinate males sired 44.0% (63/143) of chicks (Santos et al. 2015a).

### Measuring paternal care

We identified members and characterized the social group composition by conducting thorough behavioural observations within each territory (further details of behavioural work in Santos and Nakagawa 2013). The dominance status of co-breeding males was inferred from behavioural observations of displacement at feeding sites, singing perches and from the vicinity of the group's female. We recorded nestling provisioning behaviour using GoPro Hero HD cameras with 16 GB memory cards. Video cameras were set at 10 to 15 cm from the nest rim focusing into the nest cup. We filmed nests on two consecutive days (nestlings age: 8 and 9 days). Chick age had very little, and non-significant, effect on paternal care, thus it was not included in the final analyses (β_(slope of chick age)_ = 0.016, 95% CI: −0.089 to 0.129). We chose to film the nests at this stage because this is the peak provisioning period of dunnocks (Hatchwell and Davies 1990). These recordings were also used to ascertain social group composition. We quantified male parental care as the number of visits and the number of feeds during the segment of the recordings encompassed between the first provisioning visit of any adult and finishing at the end of the recorded file (defined as effective observational time, Nakagawa et al. 2007; mean effective observational time = 2.10 hours (SD: 0.573), *n =* 158 focal observations from 77 different nests).

### Statistics

We analysed our paternal care data using Bayesian mixed-effects models (BMM), with Poisson error distribution within the package *MCMCglmm* (Hadfield 2010) for *R* version 3.3.0 (R Core Team 2016). Because we have reasons to expect the number of visits and the number of feeds to be proportional to the length of the observation period, we included the natural logarithm of the effective observation time as an offset in the exposure Poisson BMMs, and fixed its coefficient to 1 (Snijders and Bosker 2011; see Supporting Information for more details). By using the offset term, we avoided having to convert the raw count data into a ratio, thus making interpretation of the results more convenient. We included the type of social group composition (categorical with 2 levels: monogamous and polyandrous), and brood size (continuous; centred and scaled) as predictors.

#### Correlation between number of visits and number of feeds

First, we estimated the correlation between the number of visits and the number of feeds because these two variables are likely to be highly correlated (see Supporting Information S1 for details of bivariate-response model to estimate the correlation). We found a strong positive correlation between the number of visits and the number of feeds (posterior mean correlation: *r_between-individual_* = 0.940; 95% CI: 0.753 to 0.985; *r_within−individual_* = 0.786; 95% CI: 0.681 to 0.859). Here, we present results of models using the number of visits as a measure of paternal care.

#### Adjustment of paternal care

As we were mainly interested in the effect of the proportion of EGY on within male paternal care, we used individual centring to separate within-versus between-individual effects in our BMMs (van de Pol and Wright 2009). This approach allowed us to quantify the effect of the proportion of EGY on paternal care among sequential broods of a particular male (within-male effect; n = 17 pairs with 2 or more breeding attempts together), and to quantify the effect between males (between-male difference; n = 66 pairs).

We found evidence of within-male adjustment of care (see Result section below); monogamous males adjust their visitation rate according to the extra-pair offspring produced by their partners. Thus, we wanted to know whether this within-individual adjustment was caused by: (i) different female partners producing different numbers of extra-pair offspring; or (ii) same female partners producing different number of extra-pair offspring in successive broods with the same male. We built another model with number of visits of socially monogamous males as the response variable. In this model, we used three fixed effects: within-male within pair deviation, within-male but between pair deviation, and between males. In addition to male identity and nest identity, we included pair identity as another random effect.

Finally, to account for the multilevel nature of our data, we included male identity and nest identity as random effects in all the models. We estimated the repeatability of paternal care in order to assess individual male consistency among different breeding attempts (see Supporting Information S1 for details). We present back-transformed (original count scale) model estimates as posterior means and their 95% credible intervals (95% CI). We considered parameters with 95% CIs not spanning zero to be statistically significant.

To test whether males reduce paternal care or not when paternity decreases, we fit the proportion of extra-group young (EGY) in a brood as a predictor in our exposure Poisson BMM. We considered the effect of within-group paternity by subordinate co-breeding males on the paternal care of dominant and subordinate co-breeding males in separate models. These analyses were a more appropriate way to compare our findings with those of Burke et al. (1989), as in their study there was no evidence of extra-group paternity, but the authors found that dominant co-breeding males in socially polyandrous groups were more likely to feed young in broods in which they gained paternity. In order to provide a similar analysis to that conducted by Burke et al. (1989) on whether subordinate males were less likely to provide care when they gained no paternity, we also investigated whether there was a difference in the number of visits by subordinate males that gained paternity in a brood, versus those that did not gain any paternity. This analysis was achieved using an exposure Poisson BMM, fitting whether subordinate males gained paternity or not as a categorical predictor.

We investigated whether paternal care was beneficial to broods by estimating the effect of the relative participation by males to the total nest visitation on the proportion of young that fledged from each nest. We fit a binomial Bayesian model, in *MCMCglmm,* with the proportion of young that fledged as the response variable (we used the *cbind* function to combine the number of fledglings and number of young that failed to fledge as the response variable), and the proportion of nest visits by males (centred, scaled) (Schielzeth 2010) as a continuous predictor. We also included the type of social group composition (monogamy or polyandry) as a categorical predictor in this model.

To assess whether paternal care is costly to males, we estimated a male’s survival until the subsequent breeding season (n = 42 males; three breeding seasons) as a function of the amount of paternal effort during a breeding season. Male paternal effort was estimated as his average provisioning visitation rate to all his nests during a given breeding season. We used a Cormack-Jolly-Seber survival model based on our mark-recapture data to estimate the effect of paternal effort in a given breeding season on his probability to survive until the next breeding season. We included the average number of visits per breeding season of the male as a covariate in the model and also the identity of the breeding season in order to control for any year effect that might cause biases in survival. Survival was estimated in the program *MARK* (White and Burnham 1999).

## Results

### General results

In total, we obtained 357 hours of paternal care footage, recorded from 66 unique nesting attempts that had paternity data available, attended by 42 and 36 unique males and females, respectively. Socially polyandrous groups accounted for 60.6% (40/66) of nesting attempts, while monogamous pairs tended the remaining 39.4% (26/66). Male number of visits increased significantly with brood size (β_(slope of brood size)_ = 0.247, 95% CI: 0.152 to 0.334). We found that co-breeding males in social polyandry made on average 0.644 times significantly less (β_(slope of polyandrous mating system)_ = −0.439, 95% CI: −0.681 to – 0.197; Figure 1) nest visits than monogamous males after controlling for the length of the observation period and brood size. Within- and between-season repeatabilities of male number of visits were significantly different from 0, and moderate to high (between seasons repeatability of paternal care = 0.794; Figure 2; see Table S1 in the Supporting Information for detailed results), which indicates that males are likely to maintain their nest visitation rates among different breeding attempts. On average, monogamous males had lower between-season repeatabilities (between seasons repeatability of paternal care = 0.452) than co-breeding males (between seasons repeatability of paternal care = 0.791), but there was substantial overlap of 95% CIs.

**Fig. 1.**
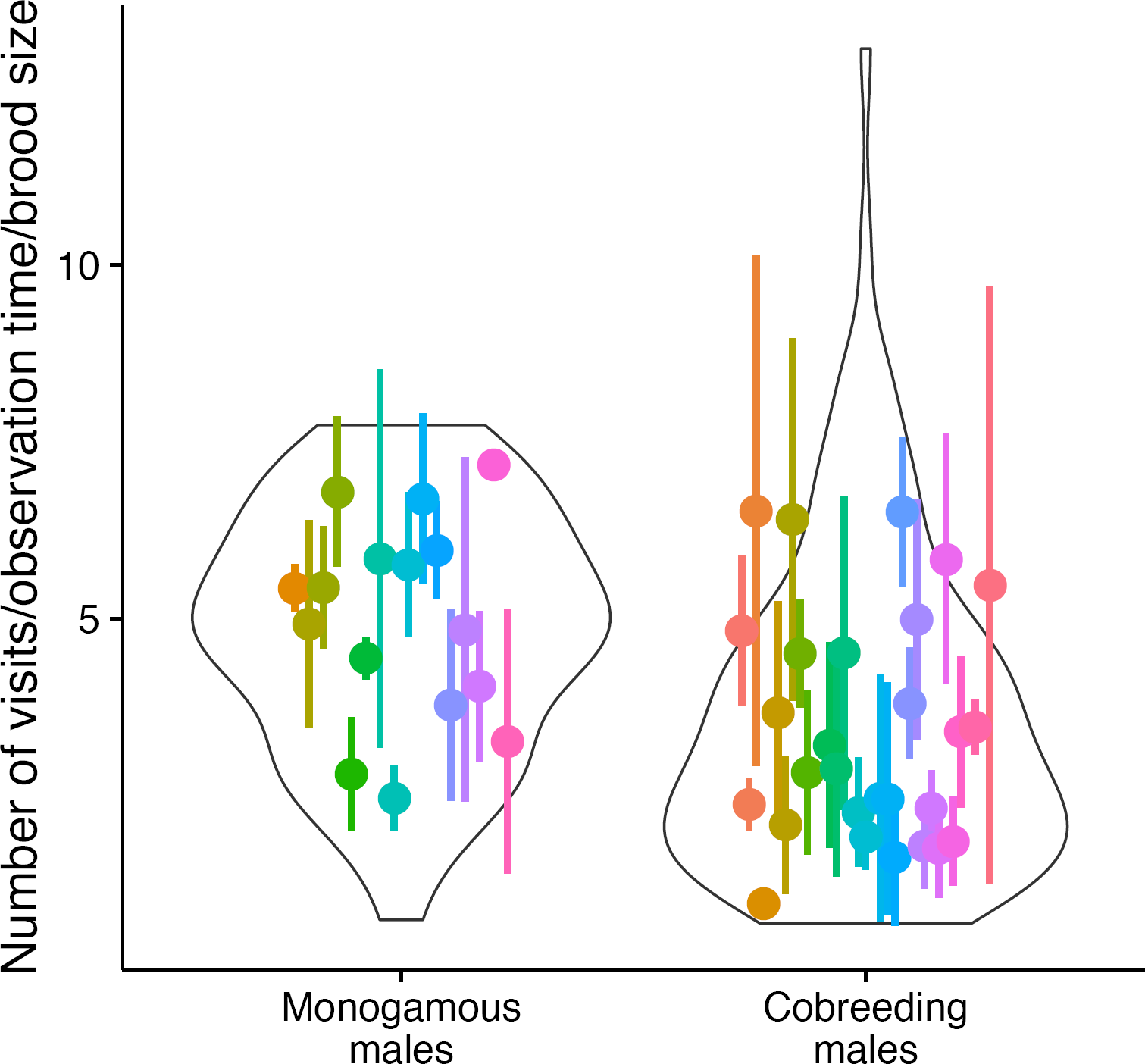
Number of nest visits by the duration of the parental care observation period made by monogamous, and co-breeding males in social polyandry (the y-axis is presented for visualization purpose; we used raw count data in our statistical models). Dots represent the mean number of visits per observation for each male (vertical bars are ± 1 SD; each male is coloured with a different shade). The envelopes show the density of the data for each male mating category.

**Fig. 2.**
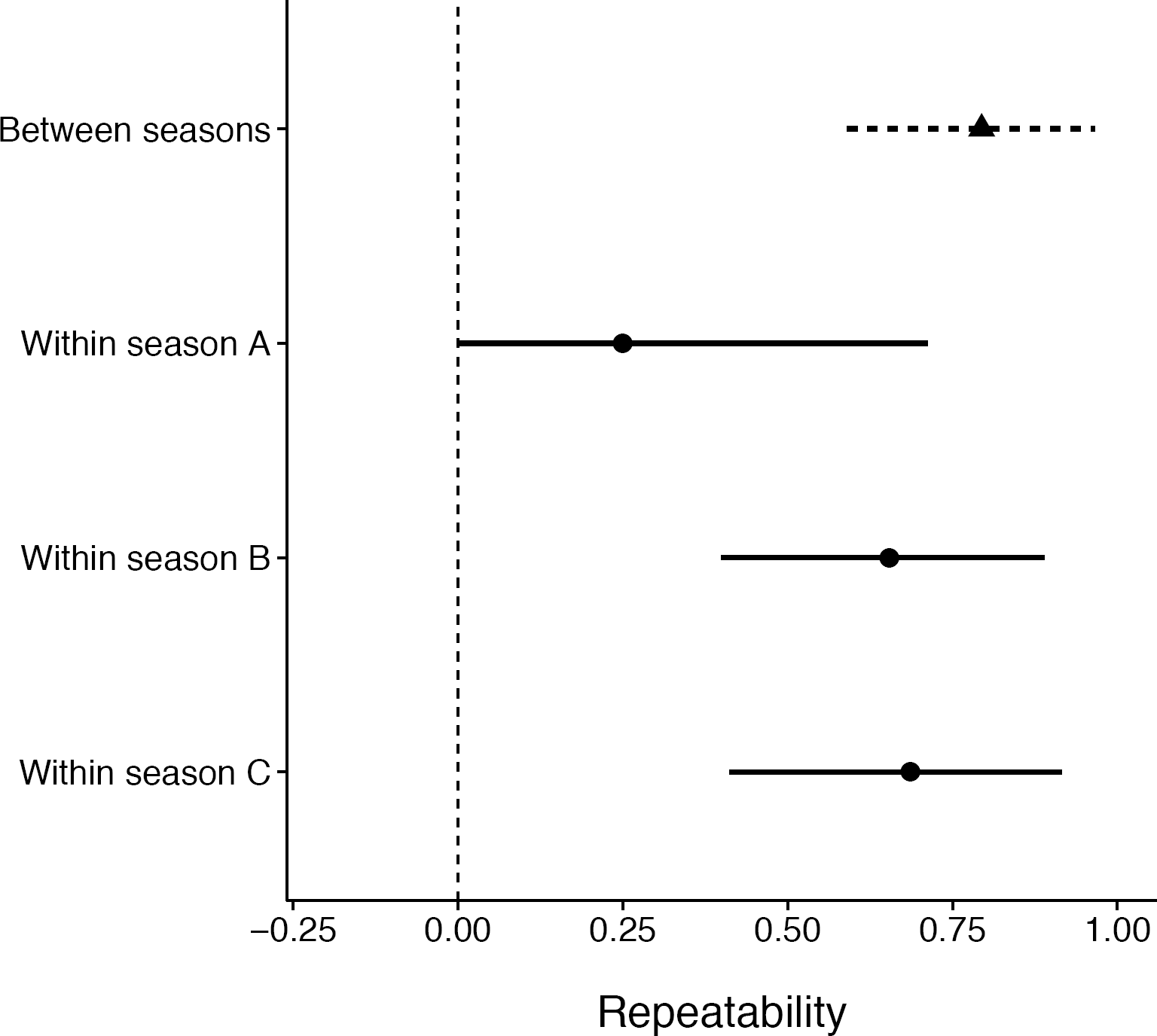
Within- and between-season repeatability estimates of the number of visits to a nest by male dunnocks. Point estimates represent posterior mean repeatabilities and horizontal bars are the 95% credible intervals. Estimates with 95% credible intervals that do not touch or overlap the vertical dashed line (0) are significantly repeatable. Season A: 2009-2010 (n = 28); season B: 2010-2011 (n = 79); and season C: 2011-2012 (n = 51).

### Paternal care and paternity

We found statistically significant evidence of a within-male effect, *i.e.* a negative relationship between the proportion of extra-pair young in a brood and a male’s number of visits, but only for socially monogamous males (monogamy: β_(within-male effect)_ = −0.732, 95% CI: −1.510 to −0.047; Figure 3). There was little evidence that co-breeding males in socially polyandrous groups changed their number of visits with changes in extra-group paternity (polyandry: β_(within-male effect)_ = 0.592, 95% CI: −0.368 to 1.448; Figure 3). We found little evidence of a between-male effect of paternity on the number of visits when examining the relationship across all pair broods (monogamy: β_(between-male effect)_ = 0.194, 95% CI: −0.531 to 0.931; polyandry: β(between-male effect) = −0.121, 95% CI: −0.943 to 0.760).

**Fig. 3.**
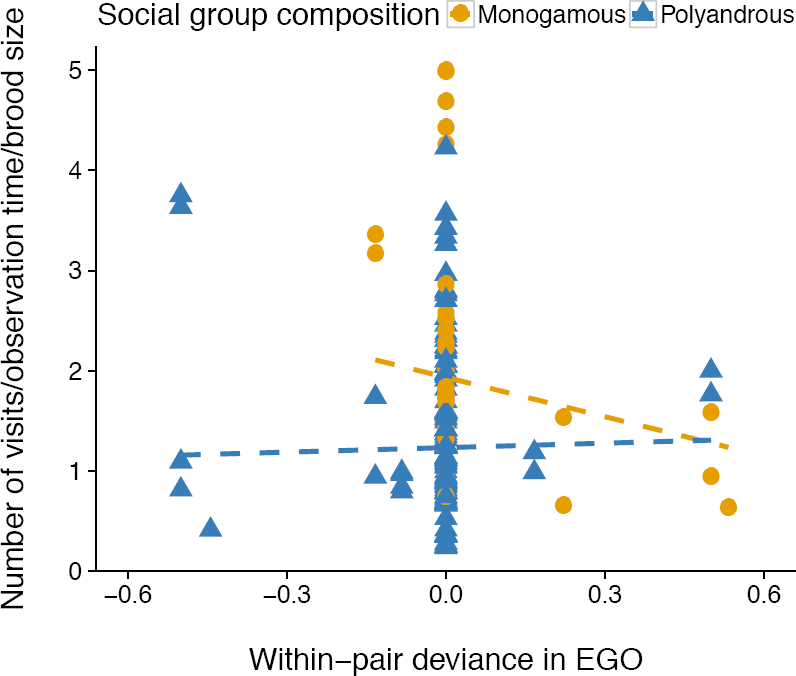
Relationship between the within-pair deviance in the proportion of extra-group offspring (EGO) in broods and male number of visits to a nest. All male-female pairs that did not show variance between breeding attempts in the proportion of extra-group offspring are spread vertically at *x* = 0. Orange and blue symbols and lines represent males in socially monogamous (n = 46 observations) and socially polyandrous (n = 112 observations) groups, respectively. Lines represent predicted results from Poisson Bayesian mixed model with *Male identity* and *Nest identity* as random effects (posterior means and 95% CI): 0.172 (0.082 to 0.293) and 0.027 (0.0003 to 0.073), respectively.

We found that individual monogamous males adjusted their paternal care within-pairs (W_mp_; monogamy: β_(within male within-pair effect)_ = −1.169, 95% CI: −2.454 to −0.023). These males reduced their visitation rate when there were more extra-pair offspring in nests of the same female partner. The estimates for the parameter within males between-pairs was negative, but the 95% CI did span 0 (W_m_B_p_; monogamy: β_(within male between-pair effect)_ =−0.665, 95% CI: −1.735 to 0.361).

In socially polyandrous groups, specifically, we found little evidence of an effect of the proportion of young sired by subordinate males on the number of visits by dominant males (β_(within-pair effect)_ = −0.346, 95% CI: −0.836 to 0.202; β_(between-pair effect)_ = 0.054, 95% CI: −0.924 to 0.985). Additionally, the proportion of young sired by subordinate males had little effect on their own number of nest visits in socially polyandrous groups (β_(within-pair effect)_ = −0.177, 95% CI: −0.921 to 0.589; β_(between-pair effect)_ = −0.674, 95% CI: −3.475 to 2.186). Finally, there was little evidence of a difference in the number of nest visits by subordinate co-breeding males in social polyandry when they gained paternity or not in a brood (β_(intercept)_ = 0.864, 95% CI: 0.509 to 1.291; β_(gained paternity)_ = −0.085, 95% CI: −0.532 to 0.381).

### Paternal care: its benefits to offspring and its costs to paternal males

We found evidence that increments in male relative participation to total nest visitation increased fledging success in monogamous broods (β_(scaled proportion male nest visits)_ = 1.942, 95% CI: 0.139 to 3.846; Figure 4), but not in polyandrous broods (β_(scaled proportion male nest visits)_ = 0.526, 95% CI: −0.572 to 1.844). There was little evidence that a male’s paternal effort influenced his own survival until the next breeding season (mean male survival = 0.892, 95% CI: 0.776 to 0.990; β_(paternal effort)_ = −0.146, 95% CI: −1.069 to 0.796).

**Fig. 4.**
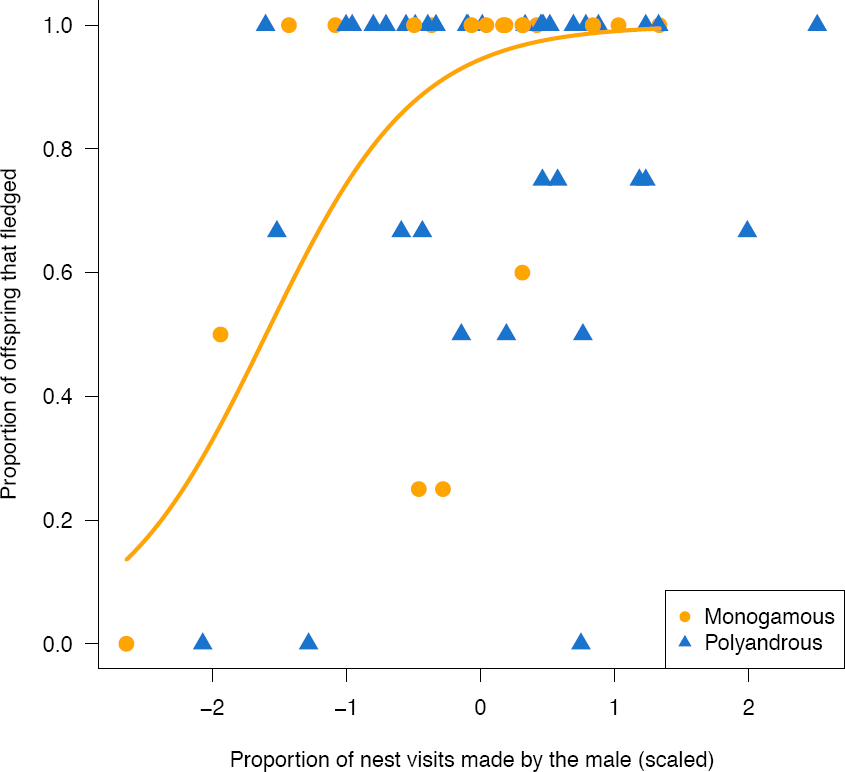
Relationship between the proportion of male visits to a nest (relative participation to total nest visitation; scaled and centred) and the proportion of young that fledged. Orange and blue symbols represent males in socially monogamous (n = 19 observations) and socially polyandrous (n = 37 observations) groups, respectively. The orange line represents predicted results from a binomial Bayesian model for monogamous males.

## Discussion

In this study, we tested whether dunnock males reduce their paternal care *(i.e*., certainty of paternity hypothesis) or not when their paternity decreases. We used a dataset with repeated observations and genetic paternity in order to address this question. Our results corroborate the certainty of paternity hypothesis, but only partially. We found evidence that socially monogamous males reduced paternal care when they lost paternity to extra-pair sires. Nevertheless, such an effect only occurred when investigating the certainty of paternity hypothesis between broods of the same breeding pair (within-male within-pair effect). We found little evidence of a between-pair effect. Below we discuss some of the implications of our findings.

Our results indicate that when within-male paternal care adjustment occurs – in socially monogamous males −, it is likely in response to changes in the extra-pair behaviour of their female partners. Which in turn suggests that socially monogamous males may be able to use information to determine how much to invest in a brood. Our findings are in contrast with predictions from a sealed-bid model (Houston and Davies 1985), in which paternal care levels would be optimized over evolutionary time. Yet, we did not find evidence that co-breeding males in socially polyandrous groups adjust their levels of care. This evidence from co-breeding males is in line with predictions from a sealed-bid model. Combined, the findings of adjustment of paternal care in socially monogamous males, and lack of adjustment in co-breeding males – in the same population – provide an interesting case about the organization of parental care with regards to behavioural flexibility. Our study indicates that the ability to adjust paternal provisioning behaviour is modulated by the breeding status of male dunnocks (see Holtmann et al. 2015 for more details about breeding status). Interestingly, our observations lead to a situation analogous to the “negotiation continuum” over the supply of provisioning behaviour (Hinde and Kilner 2007) in within-population context.

In addition to the within-male within-pair adjustment of paternal care exhibited by socially monogamous males, we found that these males’ relative participation in nest visitation has an important effect on fledging success. The combination of these two findings indicates that when socially monogamous females engage in extra-pair behaviour, they put their own broods at risk. Recently, Schroeder et al. (2016) found that male house sparrows adjust their levels of paternal care in relation to cuckoldry. However, differently from our study, Schroeder et al. (2016) found that the adjustment occurred when males changed partners that consistently differed in their levels of cuckoldry. Yet, what both studies have in common is evidence that males adjust care when cuckold, leading to costs to females (this study). Our findings, along with studies that have shown that females suffer indirect costs when producing extra-pair offspring (Hsu et al. 2015) call for further theoretical investigations of female infidelity.

Interestingly, our results do not bear similarities to another dunnock study. Davies et al. (1992) did not find evidence that the experimental removal (and subsequent loss of paternity to extra-pair males) of monogamous males from their territories during the mating period—before egg-laying—altered their provisioning behaviour. A simple explanation for the divergence between the results could be the fact that only two monogamous males were experimentally removed in Davies et al. (1992) during the period before laying. It is likely that such a small sample size would not be sufficient to detect an effect. Moreover, as we have shown in our study, the reduction in paternal care was only observed within-males within-pairs, thus, Davies et al. (1992) would not have been able to detect the effect under their experimental design.

Contrary to Burke et al. (1989), which reported that polyandrous males that gained paternity in their broods (dominant vs. subordinate males) were more likely to feed offspring than males that did not gain paternity, we found little evidence that polyandrous males adjusted their levels of care in relation to loss of paternity to both subordinate and extra-group males. Furthermore, the repeatability of paternal care was high among co-breeding males in socially polyandrous groups, indicating that males consistently provided similar levels of care in different breeding attempts. Given the previous findings and the biology of dunnocks, we found our results to be unexpected. We could simply attribute the discrepancy to differences in demography, and ecology between the two populations (our study vs. the Cambridge study). Nevertheless, we expected that co-breeding male dunnocks would be able to assess their risk of being cuckolded (i.e. having information on female promiscuity), because of the high levels of mate guarding in this species (see Davies 1992). Possibly, males have information about the risk of being cuckolded, but lack the ability to recognize their own offspring in a brood. Thus, adjustment of paternal care could be maladaptive, because males would reduce care to their own young, thus directly decreasing their own fitness. Nevertheless, our results of lack of adjustment and high repeatability among co-breeding males are consistent with the prediction of the sealed-bid hypothesis (Houston and Davies 1985) that ‘negotiation’ over how much care to provide occurs over evolutionary time (Houston and Davies 1985) (e.g., Schwagmeyer et al. 2002; Nakagawa et al. 2007). Under this scenario, changes in paternity between breeding attempts of a male should not influence how much care he provides.

Similarly to dunnocks, in the reed bunting, *Emberiza schoeniclus,* an early study found evidence for the certainty of paternity hypothesis (Dixon et al. 1994), whereas a more recent investigation did not (Bouwman 2005). In the latest reed bunting study (Bouwman 2005), the authors discussed three assumptions from theoretical models that need to be met in order for males to adjust care in relation to paternity. Succinctly (refer to Bouwman et al. 2005 for a detailed discussion), these assumptions are that: i) levels of paternity need to vary between breeding attempts of a male; ii) males should be able to assess their share of paternity through behavioural cues (e.g., absence of female during her fertile period or number of intruding males); and iii) the benefits of reducing care should outweigh the costs. The first assumption is not contentious throughout most studies, as levels of paternity vary considerably among broods (reviewed in Griffith et al. 2002). However, evidence supporting the other two assumptions is debatable. Our findings suggest that under some circumstances (social monogamy, within-pair effect), males adjust their paternal care when paternity is lost to extra-pair sires. These results indicate that these males may use cues provided by females to adjust the amount of care provided. However, it is worth noting that the within-pair reduction of care could also be triggered by other factors associated with the identity of individual females, such as their breeding condition, which could affect her share of parental care towards the brood. Disentangling the effects of factors associated with female identity from variation in extra-pair paternity would yield interesting insights into the mechanisms that cause males to reduce their paternal care. The third assumption that males should benefit from reducing their care can also be questioned. Our results suggest that males do not suffer mortality costs of providing paternal care. This finding is in agreement with a recent meta-analysis that has shown that in birds, males do not gain benefits in terms of their own survival to the next breeding season when paternal effort is experimentally reduced (Santos and Nakagawa 2012; but see Magrath and Komdeur 2003 for a discussion of benefits in terms of additional mating opportunities).

In conclusion, our results show that males are able to adjust their paternal contribution, but only under specific conditions. Monogamous males adjust their paternal investment within-pairs, and this may have important fitness consequences to the extra-pair offspring, as the amount of paternal care provided is associated with fledging success. Co-breeding males in socially polyandrous groups did not adjust their levels of paternal care according to variation in paternity. Taken together, our results raise the question of what causes socially monogamous—but not co-breeding males in social polyandry—to adjust paternal care when paternity changes when paired to the same female. Finally, females in social monogamy that engage in extra-pair behaviour face important fitness costs, as their partners’ reduction in care has negative consequences to nestling fledging success.

## Acknowledgements

We thank the Dunedin Botanic Garden, its staff, especially Barbara Wheeler, for allowing us to conduct our research on their grounds; Delphine Scheck, Patrick Crowe, Sam van der Horst, Luana L. S. Santos, and Sophie Gibson for field assistance; Luana L. S. Santos, Losia Lagisz and Karin Ludwig for their valuable help with laboratory analyses; Lonneke Kamphues and Sophie Gibson for helping to watch the parental care videos; Deborah Dawson, Andy Krupa, Daichi Saito and Isao Nishiumi for providing primer aliquots; Gustavo Requena, and Diogo Melo for thoughtful comments on an earlier version of this manuscript. ESAS received a University of Otago postgraduate scholarship, SN held a Marsden grant (UOO-0812) and a Rutherford Discovery Fellowship, and ESAS and SN received an Association for the Study of Animal Behaviour research grant.

## Supporting information

**Supporting information S1** Supplementary methods, tables and figures.

## Data accessibility

Data used in this manuscript will be made available in the figshare repository upon acceptance of the paper.

## Author contributions

ESAS conceived the study; ESAS and SN designed the study; ESAS collected the data, conducted laboratory analyses and analysed the data; ESAS and SN wrote and edited the manuscript.

